# A Pre-computed Probabilistic Molecular Search Engine for Tandem Mass Spectrometry Proteomics

**DOI:** 10.1101/2020.02.06.937870

**Authors:** Jeff Jones

## Abstract

Mass spectrometry methods of peptide identification involve comparing observed tandem spectra with *in-silico* derived spectrum models. Presented here is a proteomics search engine that offers a new variation of the standard approach, with improved results. The proposed method employs information theory and probabilistic information retrieval on a pre-computed and indexed fragmentation database generating a peptide-to-spectrum match (PSM) score modeled on fragment ion frequency. As a result, the direct application of modern document mining, allows for treating the collection of peptides as a corpus and corresponding fragment ions as indexable words, leveraging ready-built search engines and common predefined ranking algorithms. Fast and accurate PSM matches are achieved yielding a 5-10% higher rate of peptide identities than current database mining methods. Immediate applications of this search engine are aimed at identifying peptides from large sequence databases consisting of homologous proteins with minor sequence variations, such as genetic variation expected in the human population.

## Introduction

Advances in mass spectrometry (MS) have propelled proteomics toward the forefront of translational biology, with the possibility of exceeding the breadth and complexity of large-scale whole genome analysis. However, interpretation of proteomics data remains challenging due to the vast number of predicted protein sequences possible from genetic translation, splice variation, and the myriad combinations of potential post translational modifications (PTMs). Hence, common to many liquid-chromatography coupled mass spectrometry (LCMS) proteomic surveys are large numbers of unidentified spectra which continue to receive attention and research interest [1, 2]. Current use of tandem MS proteomic search engines compare observed spectra with *in-silico* modeled spectra, calculating correlation values by probabilistic methods [3, 4] or by inferred scores based on applications of signal processing and information theory [5]. Published examples of proteomic search algorithms compare each observed spectrum in a pairwise fashion with all *in-silico* spectra, ranking the comparisons by score [6]. From a collection of all rank-one peptide-to-spectrum-matches (PSMs) a false discovery rate (FDR) can be estimated [7-12] using one of two distinct approaches to define a null distribution: the well-referenced target-decoy approach, which utilizes a null distribution derived from comparisons with a database of randomized peptide sequences [8, 10, 13, 14], and a more recent decoy-free method using a null distribution comprised of alternative target PSMs [15].

While the use of FDR estimations is absolutely prudent for MS-based proteomics research, there are potential hidden issues associated with randomized decoy sequences employed as the estimated null distribution for FDR calculations [15-19]. Additionally, these effects are likely to be exacerbated as the size and complexity of both the target and decoy database increases. This is particularly challenging for researchers attempting to unlock previously unidentified fragmentation spectra or address the inherent protein diversity beyond the typical consensus proteome, ideally including known isoforms and splice variants that arises from genetic and translation variation, or within studies examining multiple variable post-translational modifications.

The goal is to develop a proteomic search engine for large sequence databases (possibly without limit), that can be distributed in a cloud computing environment while remaining comparable or competitive to existing proteomic mining algorithms. Here we present a new proteomic search engine, JAMSE (Just Another Molecular Search Engine), that utilizes a precalculated fragmentation index of all non-redundant proteotypic peptides to derive PSMs ranked by common and well developed text mining algorithms, thus basing the approach on both information theory [20] and probabilistic information retrieval [21, 22]. The result of this approach is a PSM score modeled on the relevance of indexed fragment ions, as opposed to completeness of the observed spectrum. This is not an entirely new concept for tandem LCMS, as the field of selected reaction monitoring exploits the unique combinations of elution time, precursor mass and a selected fragment mass [23].

For benchmark comparisons, a decoy-free null distribution is utilized for estimating the FDR and introduce the concept of an intrinsically measurable FDR by including protein sequences from an orthogonal species. The pre-computed nature of JAMSE renders a parallel search against a randomized decoy peptide sequences unfeasible, as not all pairwise target peptides are explored. Results, when compared to current approaches in proteomic search engines, indicate a more discriminating PSM correlation metric can be achieved. JAMSE is optimized for Linux, suitable for cloud distribution, although the efforts herein are designed to test the theory, demonstrations can be made available through https://www.jamse.app/.

## Methods

### Data

The yeast MS proteome (*Saccharomyces cerevisiae*) collected by Ishburn *et*.*al*.[24] and made publicly available, is used as a benchmark data set. Briefly, it consists of 8 sets of tandem liquid chromatography (LC) MS injections, derived from a series of orthogonal LC separations coupled to tandem MS experiments designed to comprehensively probe the yeast proteome.

### Databases

Protein sequence databases for JAMSE and Comet searches are built from subsets of the UniProtKB repository [25] containing known proteins for *Saccharomyces sp*., along with those for *Arabidopsis sp*. The inclusion of *Arabidopsis sp*. proteins is used as an intrinsic decoy to evaluate the competing methods of FDR estimation – target-decoy and decoy-free.

Enumerated in Table 1 are two distinct databases. The first, built from the UniProt consensus repository (referred to as simply UniProt herein) consisted of 7,980 *Saccharomyces sp*. proteins and 15,878 *Arabidopsis sp*. proteins, totaling 23,858 proteins with 11.5 million unique peptides. The second, built from combined UniRef100 [26] and TrEMBL [25] databases (designated UniRef herein), consisted of 58,829 *Saccharomyces sp*. and 120,863 *Arabidopsis sp*. proteins, totaling 398,119 proteins with 36.8 million unique peptides. Complimentary decoy databases are built for each of the two protein databases by randomly shuffling each protein sequence.

**Table 1.** Accounting of proteins and peptides in two distinct databases computationally constructed from UniProt repositories [21]. Peptides are generated by modeling tryptic cleavage, with the variable PTM of acetylation on K and N-term.

| Designation | Source | Organism | Proteins | Unique Peptides w/ PTMs |  |
| --- | --- | --- | --- | --- | --- |
|  |  |  |  | Individual | Union |
| UniProt | UniProt<br>(Jan 2019) | Saccharomyces sp. | 7,980 | 3,701,494 | 11,493,070 |
|  |  | Arabidopsis sp. | 15,878 | 7,813,095 |  |
| UniRef | UniRef100,<br>TrEMBL<br>(Jan 2019) | Saccharomyces sp. | 58,829 | 11,352,240 | 36,757,734 |
|  |  | Arabidopsis sp. | 120,863 | 25,493,700 |  |

### Indexing and Search

The JAMSE approach to peptide sequence identification does not consider relative peak heights, only the presence of a given peak above a threshold, and therefor is not a challenger to methods attempting to apply machine learning to spectral reconstruction [27]. Rather, JAMSE exploits patterns observed in biological polymers [28] that form the fundamental characteristics of molecular mass defect [29]. Similar to the center of mass defect distributions estimated by Mitra, et.al. [30], JAMSE employs a simple algorithm to convert a precise real number m/z value to an integer value that can be indexed by any common indexing engine, essentially binning a given mass value to the nearest mass defect distribution, which we will call a fragment mass index (FMI). We therefore treat a collection of peptides as a corpus and corresponding FMIs as indexable words, allowing us to leverage ready-built text-mining search engines and common predefined ranking algorithms [21, 22, 31-33].

Pre-computed peptides and corresponding FMIs are based on the known proteome, utilizing common proteotypic rules of digestion, considering up to two missed cleavages, while generating all possible combinations of the PTM acetylation applied to the N-terminal peptide and Lysine residues. FMIs are generated with respect to *in-silico* derived fragmentation patterns, considering both *y-ion* and *b-ion* fragments along with metastable water-loss ions.

Database searching is accomplished by querying a set of observed precursor and fragment mass values, typically filtered by abundance according to similar methods [4], for a given spectrum and converted to respective FMI values. All PSMs returned are ranked by a probabilistic score derived from a common text-mining ranking function [32] as described in Equation 1. The PSM scoring model used in JAMSE is simply the sum of the inverse document frequencies (sum_idf) across all terms, which can be interpreted as the probability that a given peptide (e.g. item/document in a corpus) *D* contains the term *t* (or FMI, as it relates to here to tandem LCMS).

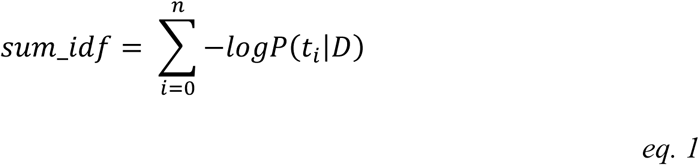

It should be noted that from this equation it is evident that rarer fragments contribute a higher score value and taken together with the overall distribution of fragment masses indicates that larger mass fragments are rarer and, therefore, contribute greater to a given PSM score. Of important note, JAMSE spectrum searching does not consider all pairwise comparisons between the observed spectrum and relevant *in-silico* spectra. Rather, much like a modern index search engine, top hits are produced via this modified method of information retrieval [21].

For this study and for comparative search parameters, all searches are limited to a 0.1 Dalton precursor neutral mass tolerance and a 0.5 Dalton fragment measured mass tolerance (i.e. mass defect width), with the top 5 hits retained per spectrum. Additionally, the Comet [34] search algorithm, specifically, the metric *xcorr*, is used the metric for direct comparison to JAMSE *sum_idf*. For computational clarity, Comet is run on a single computer utilizing 16 cores and consuming approximately 1.1GB of memory. JAMSE is distributed across two computers utilizing 8 cores each requiring approximately 1.8GB of memory per machine.

### FDR Estimations

The estimated FDR (eFDR) is computed either from the target-decoy approach as described by Keller *et*.*al*.[10], or as the decoy-free approach described by Gonnelli *et*.*al*. [15]. For the purposes of validating a given eFDR, we here introduce the concept of empirically measured FDR (mFDR). Both eFDR and mFDR methods provide means to define the null distribution and calculate the FDR q-value according to the procedure proposed by Storey *et*.*al*.[11].

#### mFDR calculation

The use of *Arabidopsis sp*. provided a means to calculate the mFDR. Within the constructed protein databases, there are approximately twice as many *Arabidopsis sp*. proteins, and if one considers false positives a random occurrence, then any *Arabidopsis sp*. peptides passing the desired cutoff represented approximately 2/3 the total number of false positives. Thus, the weighted ratio of approximately 3/2 times the number of observed *Arabidopsis sp*. peptides matched represents the mFDR. It is not necessarily suggested to include orthogonal species while searching as finding an appropriate decoy species may be challenging. To be explicit on the choice for *Arabidopsis sp*., there are only 5,643 peptide sequences shared between the two organisms given the UniProt consensus repository (approx. 5 per 10,000), and 22,434 given the UniRef100 and TrEMBL repositories (approx. 6 per 10,000). Any shared sequences identified are assigned to yeast for the purposes of further analysis.

### Evaluation of Search performance

Three distinct evaluations and comparisons are considered; the more traditional total number of PSMs per FDR, the application of estimated verses measured FDR (eFDR ∼ mFDR), and a bootstrapped estimation of the receiver operator characteristics (ROC) with corresponding area under the curve (AUC) for given search metrics. Traditional evaluation of search performance between JAMSE and Comet is based on the total number of PSMs obtained for a given eFDR q-value, with larger numbers considered more desirable. Evaluation of eFDR performance (eg. target-decoy and decoy-free) is based on the comparison to the complimentary mFDR, with a closer match to mFDR considered better. Again, it is should be noted that the decoy-free method of FDR estimation is preferred for JAMSE, as this method reduces the computational complexity of the search algorithm intended to only search sequences of natural occurrence. Bootstrap estimations of PMS discriminator metrics detailed in Table 1 was accomplished by randomly selecting a thousand PSM yielding scans, and filtered to the top peptide by corresponding metric for each scan, wherein a peptide having origin from *Saccharomyces sp*. is considered a true match and a peptide from *Arabidopsis sp*. considered a false match. Using this scheme, repeated 1000 times, allowed us to generate ROC curves and estimate the performance as a function of AUC and the true positive rate (TPR) at a false positive rate of 0.05. We choose to include the ROC as it reveals how post-processing validation and machine learning methods might exploit that curve to yield more PSMs at a given FPR.

## Results and Discussion

Search results using JAMSE demonstrate that a highly discriminant PSM score can be constructed using a model derived from information theory and probabilistic information retrieval. As detailed in Table 2, JAMSE yielded an additional 3.8% in number of PSMs at FDR 0.05. Exploring the complexity of PSM matching, as detailed in Figure 1, a select set of metrics was evaluated from the search results to compare discriminatory power. For each of the metrics a simple ROC estimation was performed, and this same procedure was applied to the Comet search results and offers a unique comparison. For instance, Figure 1a details the discriminatory power of the FMI dot-product (psm_dp) which yields a considerably higher AUC and TPR (at FDR 0.05) than Comet’s corresponding metric (Figure 2, num_matched_ions). Breaking out the dot-product components by fragment *y* and *b* ions, shows that for this particular experiment, and likely a product of the instrument and fragmentation process, *y* fragment ions impart a great contribution over their *b* ion counter parts (Figure 1a). It is perhaps surprising to discover that mass accuracy alone has little discriminatory power, (Figure 1b and Figure 2 massdiff), although upon rationalization, as researchers explore increasingly larger databases, the probability of overlapping *in-silico* spectra increases regardless of mass resolution, albeit for high-resolution measurements that probability is lower.

**Table 2.** Table detailing the proteomic search metrics evaluated for both JAMSE and Comet. Each of these metrics are evaluated by ROC in Figure 1 and Figure 2.

| Source | Metric | Description |
| --- | --- | --- |
| Comet | xcorr | cross correlation coefficient |
|  | expect | calculated expectation value |
|  | massdiff | the absolute value of the difference between the in-silico precursor mass and the observed precursor mass |
|  | num_matched_ions | the total number of fragment ions that align given the mass tolerance |
| JAMSE | pre_mma | the absolute value of the difference between the in-silico precursor mass and the observed precursor mass |
|  | psm_mma | the mean absolute value of the difference between the in-silico fragment masses and the observed fragment masses |
|  | psm_dp | the total number of fragment ions that align given the mass tolerance |
|  | psm_dp_y | the total number of y series fragment ions that align given the mass tolerance |
|  | psm_dp_b | the total number of b series fragment ions that align given the mass tolerance |
|  | sum_idf | inverse document frequency summed across each term |

**Figure 1.**
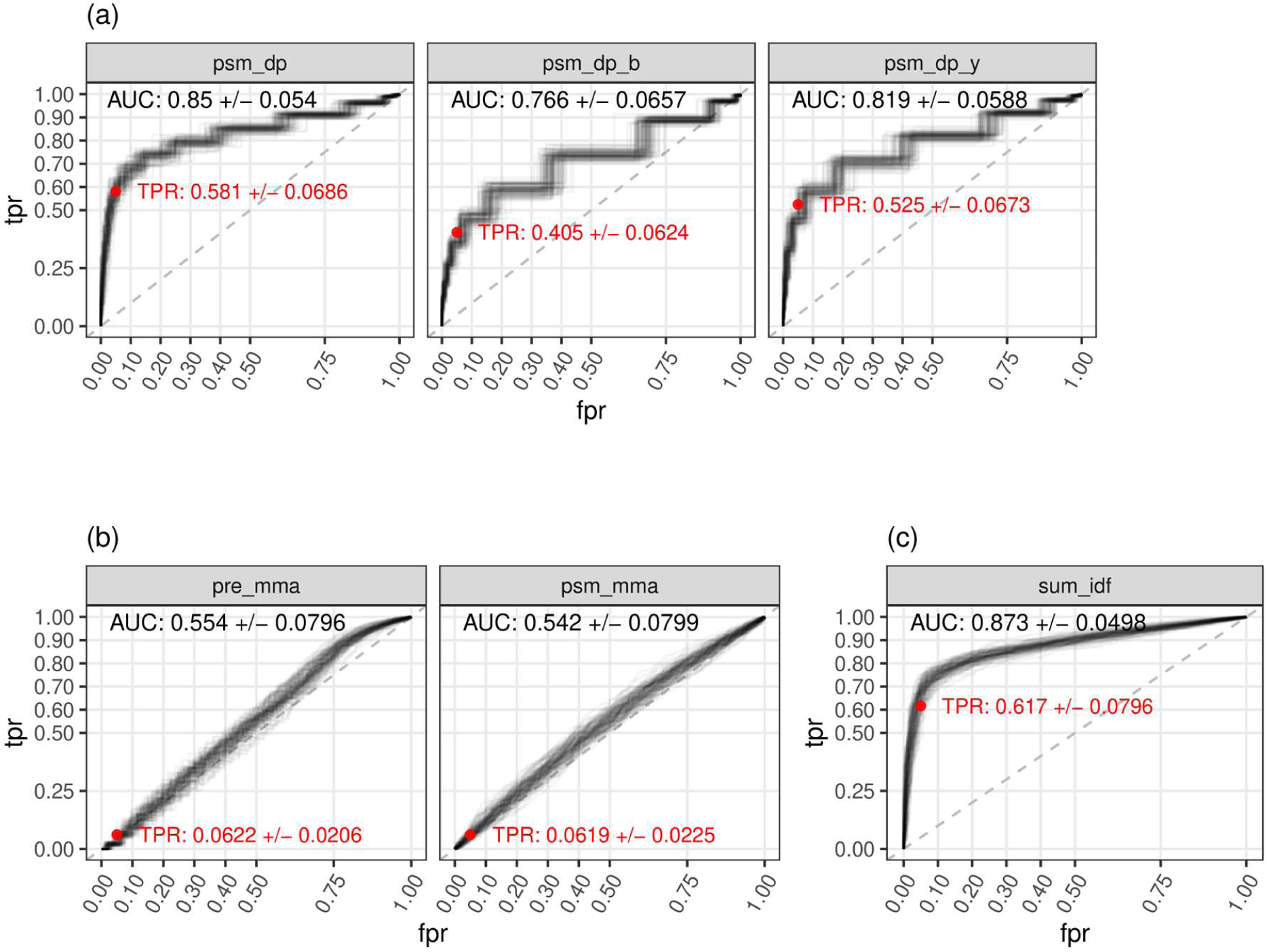
ROC plots for the discriminatory power of 100 rounds of a thousand randomly selected ms2 scans for the (a) dot-product between observed and in-silico, for all predicted FMIs (psm_dp), just the y fragment FMIs (psm_dp_y) and just the b fragment FMIs (psm_dp_b), (b) the absolute value of precursor mass measurement accuracy (pre_mma) and the absolute value of mean of fragment mass accuracy (psm_mma), (c) the ranking function selected for JAMSE, the sum of all term’s inverse document frequency (sum_idf).

**Figure 2.**
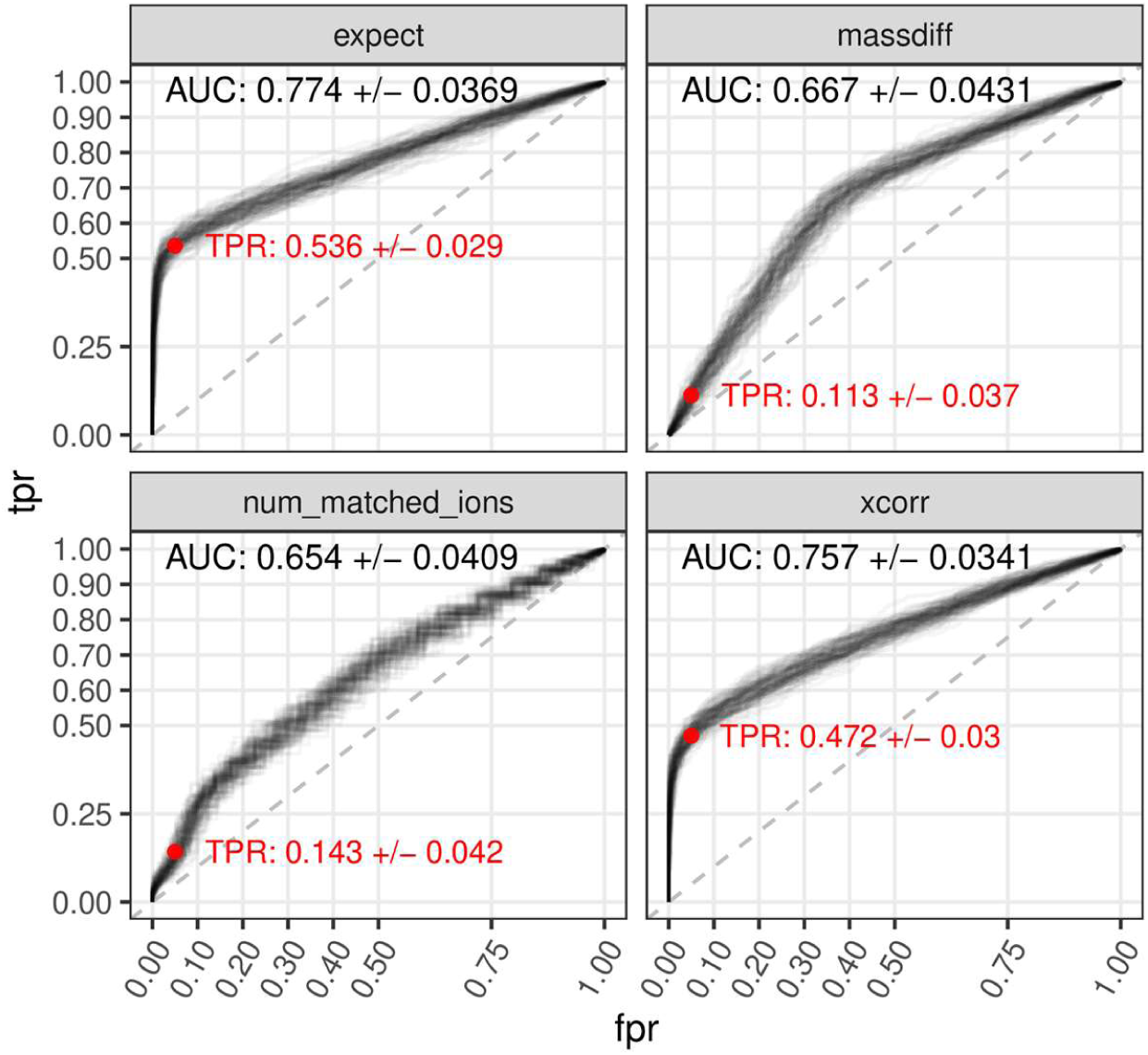
ROC plots for the discriminatory power of 100 rounds of a thousand randomly selected ms2 scans for selected the Comet metrics describing the absolute value of precursor mass measurement accuracy (massdiff), the number of matched fragment ions between observed and in-silico spectra (num_matched_ions), the cross-correlation coefficient (xcorr), and the computed expectation value (expect).

The initial goal in developing JAMSE was to explore information recall as an orthogonal means of peptide sequence identification, was to achieve comparable performance to current proteomic database mining algorithms. Figure 1c shows the bootstrapped ROC for the *sum_idf* ranking function with both a considerably higher overall AUC and TPR (at FDR 0.05) than Comet’s two corresponding metrics (Figure 2, xcorr and expect). It should be noted that current methods of scoring PSMs do not consider the rarity of given fragment ions, and instead compute a score or probability based on the distribution of observed completeness of fragmentation for all pair-wise comparisons [4-6, 34-38]. It is reasonable to assume that some sequences may be identifiable with few fragments forming a rare combination, while other peptides with a shared or common fragmentation pattern require a near complete representation; JAMSE exploits this relative uniqueness of fragmentation patterns. Additionally, it should be noted that not all experimental objectives require complete sequencing of peptides; many research objectives seek a balance between PSM accuracy and overall yield depending on experimental goals [39-41].

The use of the decoy free method for FDR estimation comes from the need to avoid additional computational resources required to search a reverse sequence database; as the size of the database grows the chance of having iso-spectral overlap increases. Additionally, there is uncertainty in the randomized and shuffled decoy creation process to generate natural sequences, after all only a small fraction of all possible sequence combinations from observation and predicted transitions have been catalogued.

Considering the UniProt database (Table 3), JAMSE yielded more PSMs than Comet for a given decoy-free eFDR value (Figure 3a, blue and red solid lines respectively), demonstrating that PSMs generated through JAMSE’s probabilistic information retrieval had higher discriminative power than did those from traditional cross-correlation. Although JAMSE found more PSMs, there is very good agreement in the peptides that are identified by both methods; PSMs that differed across the methods often contained the same peptide sequence and differed only in the PTM location (data not shown). For example, at an eFDR (decoy-free) of 0.01, Comet yielded 20,755 PSMs, while JAMSE yielded 23,943 PSMs, a substantial 15.4% gain, and a further look at the overlap in PSM assignments showed 92.6% agreement at FDR 0.01, using the smaller UniProt database (Figure 4, red line). Using the larger UniRef database, the agreement between search algorithms is 89.6% at FDR 0.01, within range of similar comparisons by others [14].

**Table 3.** Table comparing the eFDR and associated mFDR results for both Comet and JAMSE using the target-decoy and decoy-free methods. The column designated dFDR denotes the difference between mFDR and eFDR.

| Database | eFDR | Method | Algorithm/Method | PSMs | mFDR | dFDR |
| --- | --- | --- | --- | --- | --- | --- |
| UniProt | 0.01 | target-decoy | Comet | 21,404 | 0.007 | -0.003 |
|  |  | decoy-free | Comet | 20,755 | 0.005 | -0.005 |
|  |  |  | JAMSE | 23,943 | 0.011 | 0.001 |
|  | 0.05 | target-decoy | Comet | 24,639 | 0.028 | -0.022 |
|  |  | decoy-free | Comet | 25,796 | 0.045 | -0.005 |
|  |  |  | JAMSE | 26,768 | 0.041 | -0.008 |
| UniRef<br>(UniRef100<br>& TrEmbl) | 0.01 | target-decoy | Comet | 18,265 | 0.004 | -0.006 |
|  |  | decoy-free | Comet | 19,279 | 0.006 | -0.004 |
|  |  |  | JAMSE | 21,816 | 0.013 | 0.003 |
|  | 0.05 | target-decoy | Comet | 21,750 | 0.017 | -0.033 |
|  |  | decoy-free | Comet | 24,293 | 0.046 | -0.004 |
|  |  |  | JAMSE | 24,658 | 0.041 | -0.009 |

**Figure 3.**
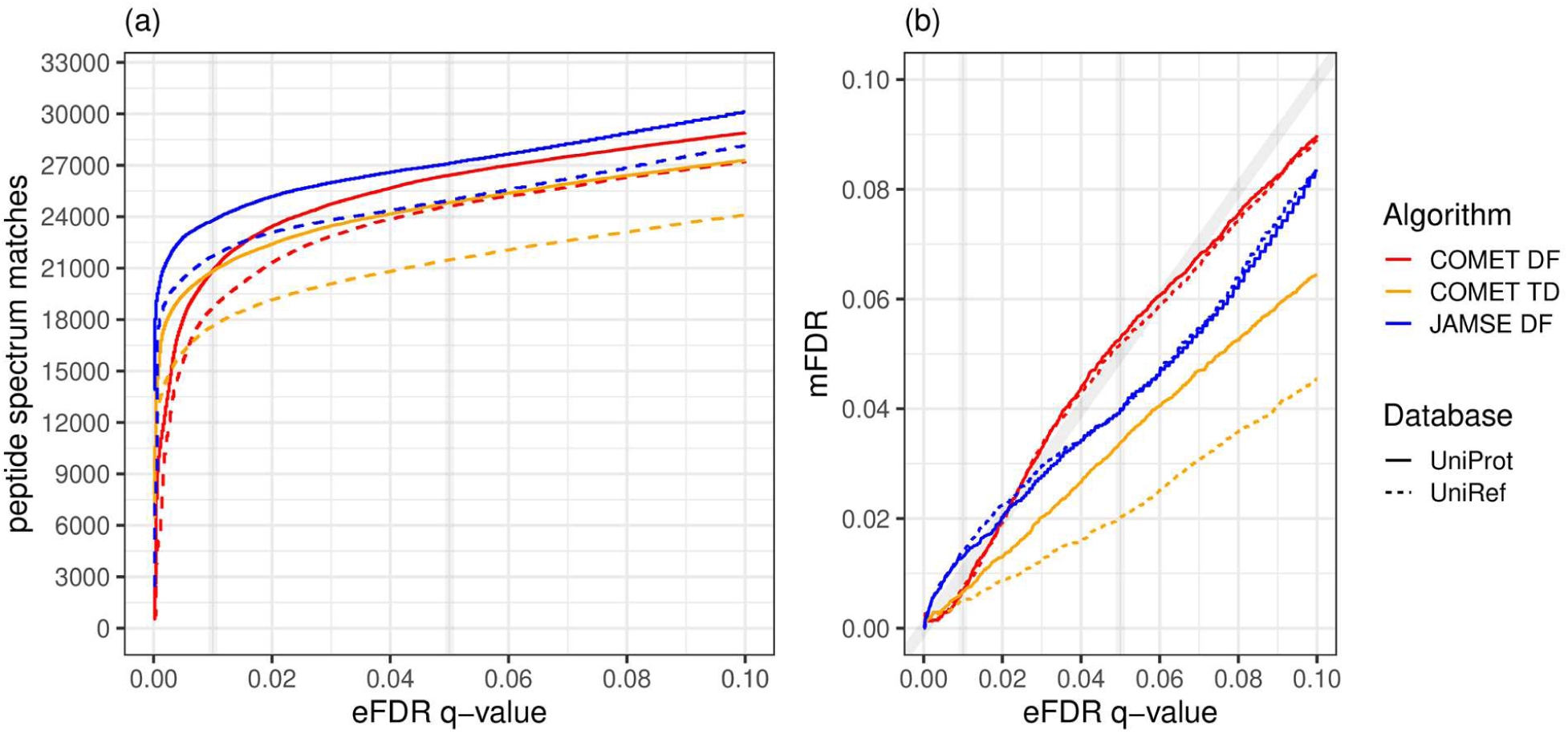
(a) Plot of PSMs recalled as a function of the FDR (based on Comet xcorr and JAMSE sum_idf) given the databased used for searching, either the conservative UniProt (solid line) consensus sequences or the combined UniRef and TrEMBL (dashed line) protein sequences. (b) Comparison between eFDR and mFDR where the solid grey line represents the function x=y, where the method of FDR estimation (eFDR) matches that of the species-specific mFDR.

**Figure 4.**
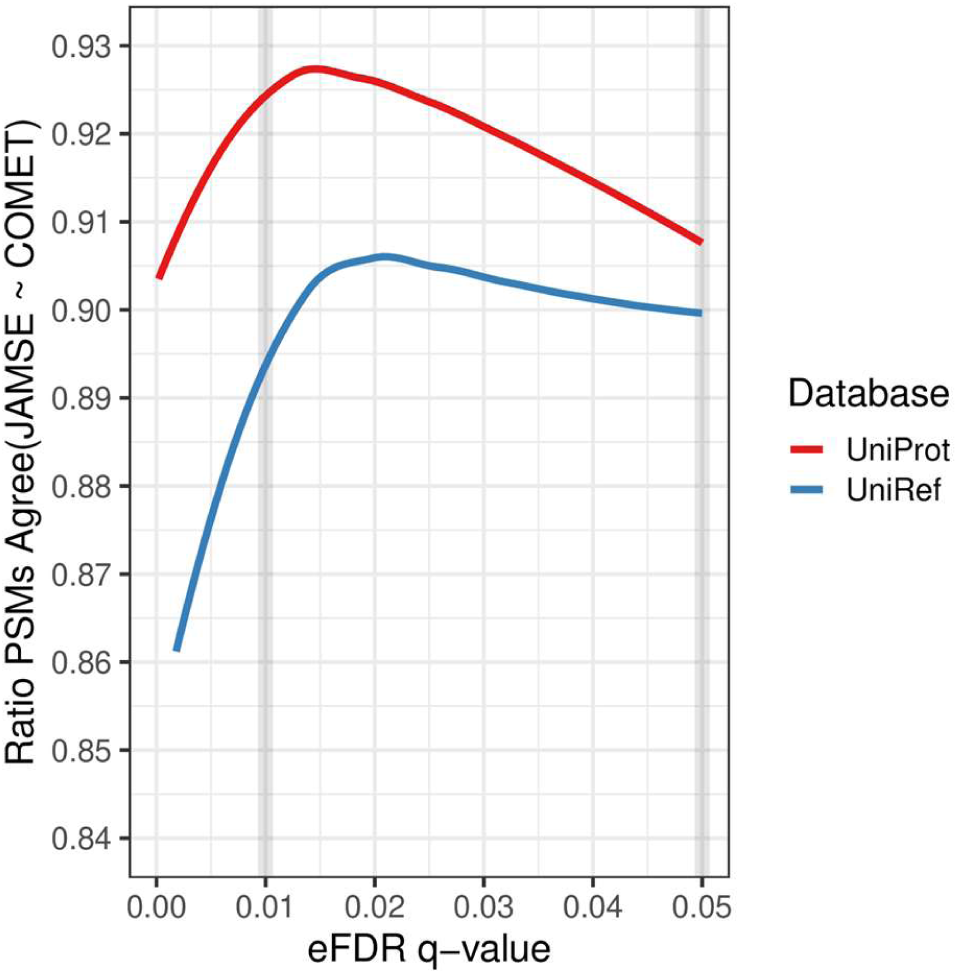
Plot of the fractional agreement between Comet and JAMSE as a function of the eFDR (decoy-free method) given the databased used for searching, either the conservative UniProt (red line) consensus sequences or the combined UniRef100 and TrEmbl (blue line).

Exploring the target-decoy eFDR for Comet (Figure 3a, orange solid line) there appears to be an early advantage, over the decoy-free method (red solid line), that diminished quickly at larger cutoffs. It is not known why this occurred, however, it is speculated that contributions from highly similar peptides sequences present in the decoy-free method are more common, as the effect is more pronounced in the larger UniRef search results. To note, JAMSE reports similar sequences as alternates with a computed Bayesian posterior probability, akin to estimating the probability of PTM location (data not shown).

When considering the larger UniRef database (Figure 3a, dashed lines) the effect of diminished returns with larger databases is evident (dashed lines lower than solid lines of same color). However, the effect is larger for the decoy-free method (orange lines) than for the target-decoy method (blue and red lines). This difference may have been due to a shift to smaller peptides in the target-decoy’s randomized protein sequence database (data not shown) further exacerbated by the highly homologous nature of the UniRef100 database. The generation of the randomized target-decoy protein sequences tends to produce 10% more peptides of sequence length 6 to 16. Consequently, nearly half of all PSMs for the target-decoy yeast dataset (52%) are between 6 and 16 residues in length (data not shown). This presents a risk in using the target-decoy approach, as FDRs may become inflated due to the proportional increase in short peptide decoy sequences relative to the target database [42].

The results for the mFDR approach (Figure 3b) are a strong commentary on the effectiveness of the decoy-free over the target-free eFDR methods. For any eFDR method, an mFDR should match, and, given the ability to compute q-values, a plot of mFDR against eFDR should follow the identity line (y = x: grey line). However, the lines depicting the relationship between mFDR and eFDR for the target-decoy approach (orange lines) indicated an overestimation of the eFDR. As above, this may have been due to the bias in peptide sequence lengths in randomized protein decoy databases and other effects noted in literature [43, 44]. On the other hand, the decoy-free approach for both Comet and JAMSE (Figure 3b, red and blue respectively) demonstrated a closer agreement between eFDR and mFDR (Table 2) with no observable difference given database size, unlike that observed for the target-decoy method.

## Conclusions

This work shows that probabilistic models in information recall can competitively and accurately identify peptides regardless of the size of the database. JAMSE provides a significant improvement for researchers attempting to address peptide sequencing in increasingly complex samples, such as the human proteome collected during clinical research and other genetically diverse sets. Peptide identification strategies have demonstrated weaknesses and therefor it is prudent to continue the development of alternatives [44] and in particular, continue to improve upon the initial PSM correlation metric, of which may enhance the post-search validation and machine learning methods. Providing a platform to expose all known sequence variants to direct comparison can augment these methods. Essentially providing an orthogonal method to unlocking the full translational potential of proteomics.

Despite JAMSE’s ability to leverage larger databases, we realize there is also a fundamental limitation to controlling the error rate, especially when larger collections of independent hypothesis are utilized [11, 45, 46]. A veritable Catch-22, increase the search space to identify rarer peptides and decrease the overall yield post FDR analysis. While there are more sophisticated methods for peptide validation, the work described herein is seen as a compliment to post-identification machine learning validation [15, 47] and methods aimed at spectral reconstruction [27]. In addition, this work is not intended to challenge precision mass measurements, rather, given the effective conversion to unit resolution via the association to fragment mass defect distributions (e.g. FMI), this method may improve the yield and accuracy from lower resolution instruments typically utilized in high-throughput clinical research.

Lastly it is hypothesized, that although not specifically demonstrated here, that JAMSE directly solves the efficiency inherent in searching a very large and homologous redundant sequence space [6, 48, 49], since JAMSE’s underlying technology leverages common indexing and search methods which can be distributed among many computer cores, or a compute cloud in parallel. Additionally, precalculated databases like JAMSE’s also introduce a path towards real-time identifications [50], since search times of less than 10 milliseconds per spectrum are easily achieved given the computational resources and database sizes used here. Future implementations could be used in tandem LCMS data-driven acquisition methods that incorporate peptide sequence identification.

## Acknowledgments

This research did not receive any specific grant from funding agencies in the public, commercial, or not-for-profit sectors.

